# Polyunsaturated fatty acid arachidonic acid increases the virulence of gut pathogen, *Yersinia enterocolitica*

**DOI:** 10.1101/2021.12.06.471175

**Authors:** Denise Chac, Kelly Crebs, Cara Yee, R. William DePaolo

## Abstract

Food-borne illnesses are a major health concern worldwide. While 1 in 6 individuals are infected in the United States yearly, there is little research into which dietary factors can alter the risk of infection. Despite evidence suggesting a correlation between obesity and enteric infection, the few reported studies focus on the role of dietary factors and the impact on host tissues and susceptibility. The direct impact of dietary constituents on the virulence of a pathogen has largely been ignored. One component of the Western diet that has been correlated with increasing inflammatory diseases is increased consumption of omega-6 polyunsaturated fatty acids such as arachidonic acid. Here, we show that arachidonic acid directly alters the pathogenicity of the food-borne pathogen *Yersinia enterocolitica*. Using *in vitro* cellular adherence assays, proteomic peptide mass fingerprint profiles and *in vivo* mouse models, we show that arachidonic acid can alter the pathogenesis of *Y. enterocolitica* by increasing proliferation and intracellular invasion. These findings have major implications in more than food safety, potentially revealing how current dietary habits may increase the virulence of food-borne pathogens.

## INTRODUCTION

Polyunsaturated fatty acids (PUFAs) play a vital role in human development and health. While deficiencies in PUFAs can lead to skin and neurological impairment, excessive PUFAs can cause inflammation.^1,2^ The omega-6 fatty acid arachidonic acid is especially of interest as it has become highly available in foods with plant derivatives, such as corn oil, and is linked to inflammatory diseases such as obesity^3,4^ and diabetes.^2,5^ Whereas studies of arachidonic acid in mice have been inconsistent in regards to its connection to colonic inflammation,^6–9^ insulin resistance,^10,11^ and obesity,^10–12^ human studies have found excessive levels of arachidonic acid among patients with obesity,^3,4^ metabolic dysregulation,^13,14^ and ulcerative colitis.^15–18^

Arachidonic acid and its derivatives such as eicosanoids are also important for antibacterial functions. Arachidonic acid is highly bactericidal against gram-positive species including *Neisseria, Pseudomonas*, and members of *Enterobacteriaceae*^19^ and reduces growth and adherence of gram-positive probiotic *Lactobacillus* species.^20^ In a recent murine study, serum arachidonic acid levels increased following *Streptococcus pneumoniae* infection and *in vitro* assays demonstrated its antimicrobial activity on the pneumococcal membrane.^21^ The effect of arachidonic acid on gram-negative pathogens is more variable. While arachidonic acid is less potent on *Escherichia coli*^19^ it can be incorporated into the bacterial membrane of *Acinetobacter baumannii* and reduce bacterial fitness and membrane integrity.^22^

Several food-borne pathogens, including *E. coli, Salmonella*, and *Yersinia enterocolitica*, are gram-negative. According to the CDC, food-borne illnesses affect 48 million individuals and have caused 128,000 hospitalizations and 3,000 deaths each year.^23^ In less developed countries, food- and water-borne diarrheal diseases have killed approximately 1.9 million people.^24^ One such food-borne pathogen is gamma-Proteobacteria *Y. enterocolitica. Y. enterocolitica* causes self-limiting gastroenteritis and mesenteric lymphadenitis in healthy individuals and can be lethal in immunocompromised patients or young children.^25^ There is also a disproportionate incidence of *Y. enterocolitica* infection found among children less than 1 year of age.^26^

During infection, *Y. enterocolitica* encounters various host environments and contains several temperature-dependent virulence factors. While at room temperature (30°C or lower), *Y. enterocolitica* is highly motile and proliferative, and expresses a chromosomal invasion factor *invasin*.^27^ Virulence factors present on a 70-kb plasmid (pYV) are expressed upon entry into the host tissue due to exposure to an increase in temperature to 37°C.^27–29^ These genes include proteins involved in colonization, effector proteins such as *Yersinia* outer proteins, *Yersinia* secreted proteins, and a type III secretion system used to inject host cells with these effectors.

The effect of arachidonic acid on *Y. enterocolitica* has not yet been established despite its potential for exposure to arachidonic acid in meats^30^ and in the gut lumen.^31–33^ There is a lack of studies investigating foodborne bacterial pathogens in the presence of PUFAs, both dietary and host derived. Studies of other dietary fats find saturated fatty acid, lauric acid, to inhibit *Y. enterocolitica* growth but not myristic and palmitic acid.^34^ Another study found oregano and nutmeg essential oils ineffective against bacterial colonization in chicken.^35^

In the present study, we found that PUFAs differentially affect the growth of *Y. enterocolitica*. Exposure to arachidonic acid increases the proliferation of *Y. enterocolitica* at both room and host temperatures. At 37°C, arachidonic acid increases its ability to invade into host cells and alters disease pathogenesis. Under normal conditions, *Y. enterocolitica* exhibits different virulence states depending on mammalian cell presence; when arachidonic acid is added, *Y. enterocolitica* becomes hyper-virulent regardless of temperature and presence of mammalian cells. These results highlight the necessity of investigation into how dietary fats can impact the virulence of food-borne pathogens.

## RESULTS

### Polyunsaturated fatty acids differentially affect proliferation of *Yersinia enterocolitica*

Recent studies involving PUFAs have shown both beneficial and detrimental effects on bacterial pathogens.^36,37^ However, there are currently no studies investigating the effect of omega-6 or omega-3 PUFAs on gut pathogen *Y. enterocolitica* (Ye). In this study, we supplemented bacterial growth media with different concentrations of omega-6 PUFAs [e.g., linoleic acid (LA) and arachidonic acid (AA)] and omega-3 PUFAs [e.g., alpha-linolenic acid (ALA), and eicosapentanoic acid (EPA)]. The concentrations of PUFAs tested were physiologically relevant concentrations found in animal-derived foods.^38^ We monitored the growth of PUFA-conditioned Ye at room temperature (26°C) by measuring OD600.

Treatment with omega-3 and omega-6 PUFAs had variable effects on the proliferation of Ye at room temperature. Cultures treated with LA, ALA, and EPA had a modest increase in proliferation and were only significant at the 500 μM concentration (**Figure 1A**). In contrast, cultures treated with AA had the largest and most significant effect on proliferation which occurred in a dose-dependent manner and seemed more sensitive to AA, as we observed significant increases at both 250 μM and 500 μM. To compare and analyze the effect across all the PUFAs, the experiment was repeated using the 500 μM concentration which had the most significant effect on Ye. We found that AA significantly increased the growth of Ye over the course of 4 hours (**Figure 1B**). EPA was the only other PUFA that had an effect at 500 μM and this occurred only at the 120 minutes, whereas ALA and LA had no effect on proliferation. Proliferation of Ye primarily occurs at 26°C. When exposed to higher temperatures within the host, energy used for proliferation and cell division is used to activate the Ye virulence plasmid (pYV), which encodes the type 3 secretion needle, as well as other factors necessary for its virulence.^39^ We then assessed whether the PUFAs had similar effects on proliferation at 37°C. Using the 500 μM concentration for all the PUFAs, we found that all four of the PUFAs tested, regardless of being an omega-3 or omega-6 PUFA, significantly increased the growth of Ye compared to vehicle treated bacteria (**Figure 1C**).

**Figure 1.**
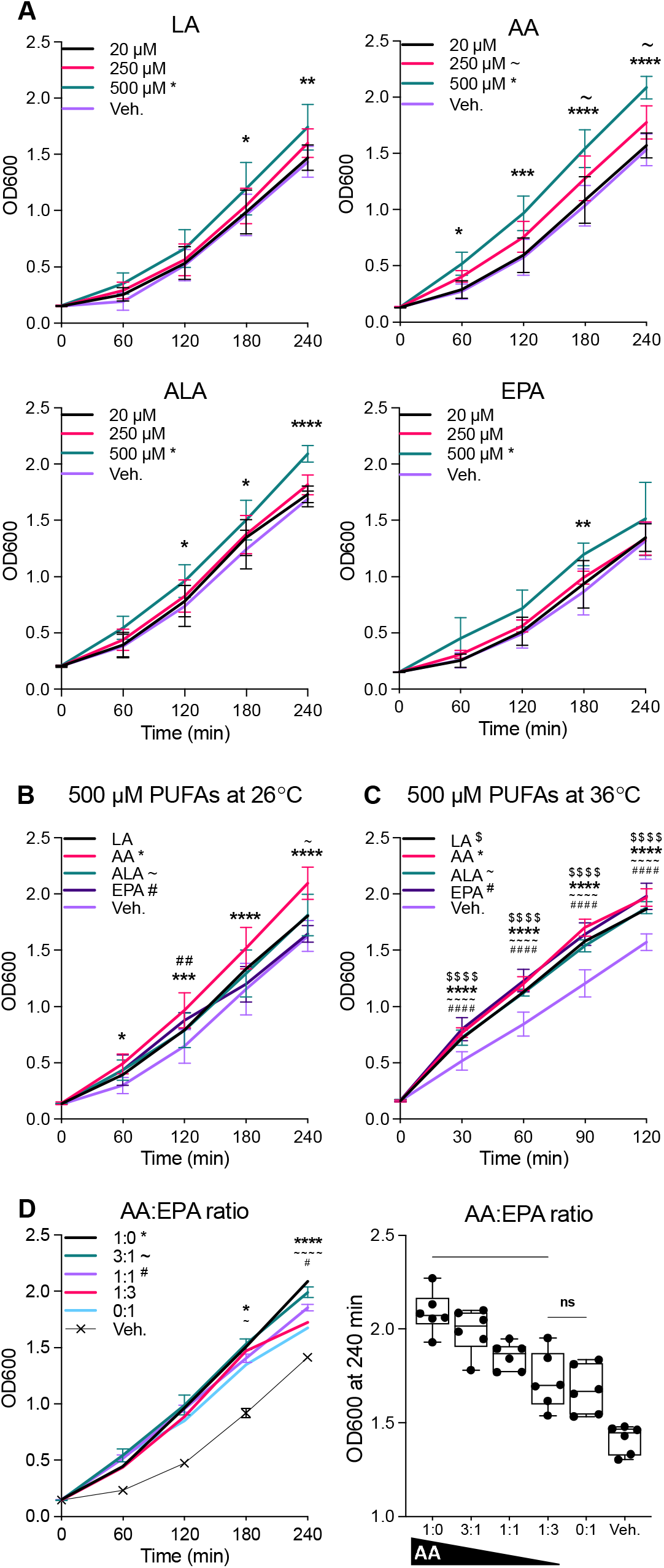
Arachidonic acid increases proliferation and intracellular invasion of *Y. enterocolitica*. **(A)** Growth curve of Ye grown with addition of 20-500 μM of linoleic acid (LA), arachidonic acid (AA), alpha-linolenic acid (ALA), and eicosapentaenoic acid (EPA). Vehicle (Veh.) control was equal volume ethanol. **(B)** Growth curve and overall percent growth of Ye over 4 hours following addition of 500 μM of PUFAs. **(C)** Growth curve and overall percent growth of Ye with ratios of AA to EPA: 500 μM AA (1:0), 250 μM AA to 250 μM EPA (1:1 ratio), 125 μM AA to 375 μM EPA (1:3 ratio), and 500 μM EPA (0:1). (A) Two-way ANOVA, multiple comparisons. Data is mean ± SEM from 3 independent studies with duplicates. * = 500 μM, ~ = 250 μM. \**P*<0.05, \*\**P* <0.01, \*\*\**P* <0.005, \*\*\*\**P* <0.001. (B, C) Kruskal-Wallis Test with Dunn’s multiple comparisons test. Data is shown as mean ± SEM from 3 independent studies with duplicates. \**P*<0.05.

Omega-3 fatty acids, such as ALA and EPA, are known to produce anti-inflammatory eicosanoids and inhibit inflammatory pathways in innate immune cells.^40^ In contrast, high dietary intake of omega-6 PUFAs is linked to inflammation and produce inflammatory eicosanoids.^41^ Both types of fatty acids compete for the same desaturation enzymes and excessive omega-6 PUFAs can inhibit the anti-inflammatory effects of omega-3 PUFAs.^41^ As the recommended dietary intake of omega-6 to −3 fatty acid ratio is 1:1 to 3:1,^42,43^ we tested whether increasing the concentration of EPA would antagonize the proliferative effects of AA. We cultured Ye with different concentrations of EPA while maintaining a constant concentration of AA. Consistent with our previous results, AA alone caused a significant increase in growth over a 4-hour period compared to vehicle alone (**Figure 1D**). When an equal amount of EPA was added to the cultures, there was a small but significant reduction in the AA-induced growth. Adding in 3 times more EPA caused a large and significant reduction in the proliferation of Ye, though not as low as the levels of the vehicle control (Figure 1D). Taken together, these data illustrate a temperature-specific effect of AA on the growth of Ye, which can be inhibited by the addition of EPA.

### Arachidonic acid induces a virulent state in *Yersinia enterocolitica*

At host temperatures of 37°C, Ye will activate its virulence plasmid and begin to produce and secrete *Yersinia* outer proteins (Yops) and *Yersinia* secreted proteins (Ysps) that are its major virulence effectors.^28,29^ As treatment with AA caused an increase in proliferation during room temperature growth, we evaluated whether there was also activation of the Ye virulence program. AA was added to Ye cultures grown at room temperature and the bacterial supernatants and cell lysates were assessed by SDS-PAGE and Coomassie blue staining for protein production. As expected, untreated Ye grown at room temperature had no staining in either the cell lysate or the supernatant (**Figure 2**). This was in contrast to the cultures grown at 37°C which showed the characteristic staining of Yops and Ysps in both the bacterial lysates and in the supernatant. While addition of the vehicle did induce a slight increase in proteins visualized by Coomassie, when AA was added to room temperature cultures, we observed a staining pattern that was much more similar to those grown at 37°C than at room temperature. These data demonstrate that at room temperature AA treatment influences proliferation and activation of the virulence program encoded by the pYV plasmid.

**Figure 2.**
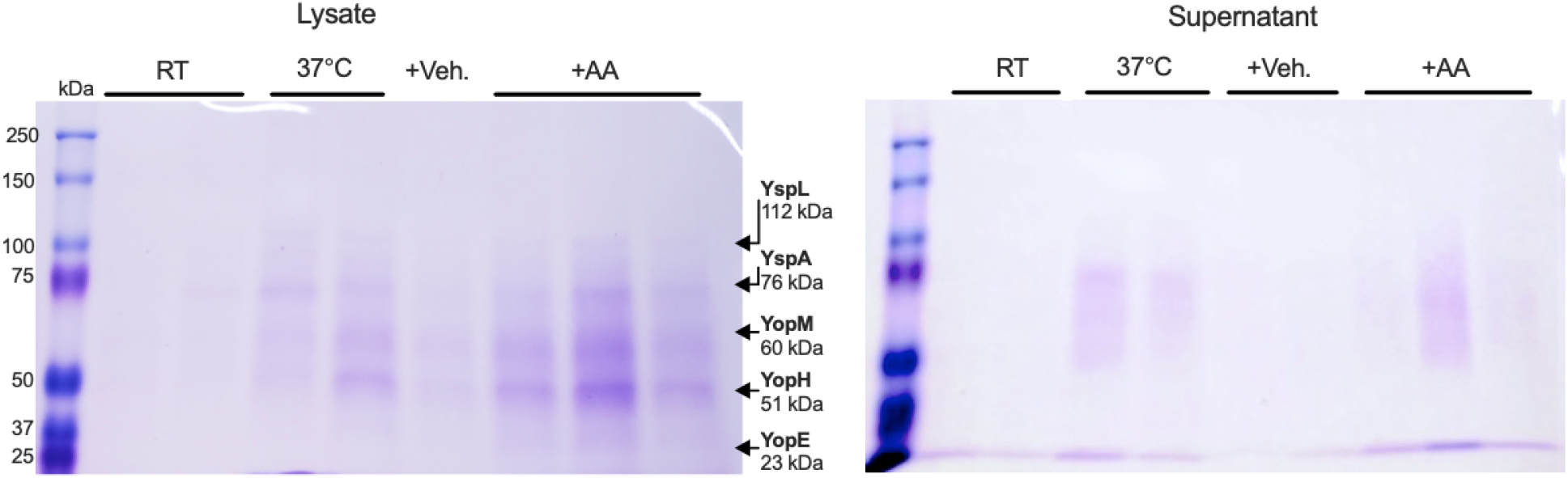
Arachidonic acid alters the virulence state of *Y. enterocolitica*. Coomassie blue staining of *Y. enterocolitica* lysates and supernatant grown at room temperature (RT), 37°C, or exposed to 500 μM AA or vehicle at RT.

### Arachidonic acid increases intracellular invasion of *Y. enterocolitica in vitro*

To determine whether AA impacts Ye colonization, we tested adherence and invasion using intestinal epithelial cells. Ye was conditioned with AA for two hours at 26°C, washed in PBS, and cultured with the human intestinal epithelial cell line, SW620. Bacteria were recovered after 1 hour to analyze the percentage of Ye adhered to cells or the SW620 cells were treated with gentamicin and bacteria was recovered an hour later to analyze intracellular Ye (**Figure 3A**). Supplementation of the cultures with AA resulted in nearly a 2-fold increase in adherent Ye and a significant increase in the amount of Ye recovered intracellularly compared to untreated and vehicle controls (**Figure 3B**). Interestingly, the addition of EPA was able to reduce the AA-induced invasion when added at a 1:3 ratio of AA to EPA (**Supplemental Figure 1**).

**Figure 2.**
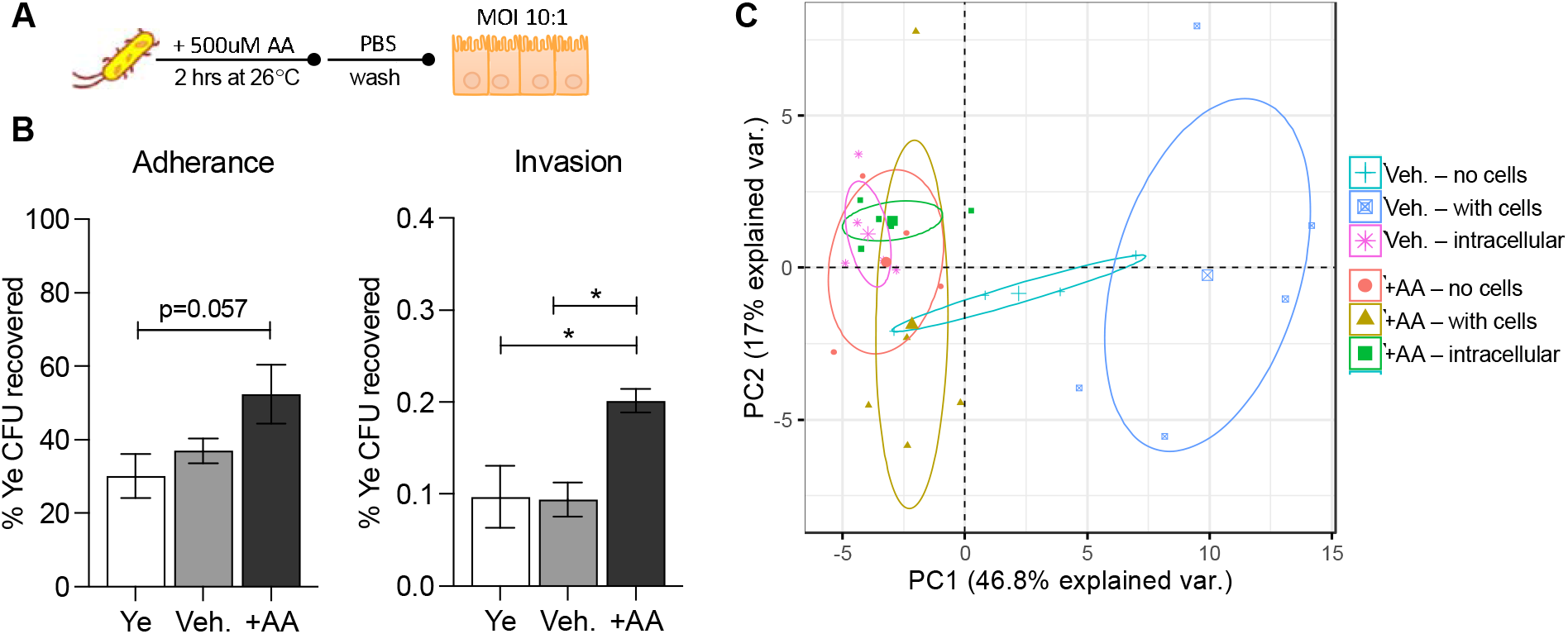
Exposure to arachidonic acid alters invasion and peptide mass fingerprint of Y. enterocolitica. **(A)** Diagram of assay using *Y. enterocolitica* exposed to 500 μM AA and then introduced to human colonic epithelial cells SW620s. **(B) (left)** Percentage of *Y. enterocolitica* CFUs recovered after 60 minutes of incubation and **(right)** percentage of *Y. enterocolitica* CFUs recovered after 60 minutes of incubation then treatment of 100 μg/ml gentamicin for 60 minutes. (B) One-way ANOVA with Tukey’s multiple comparisons test, \**P*<0.05. Data is mean ± SEM from a representative from 2 independent studies with duplicates. (C) Principal component analysis of peptide mass fingerprint profiles of *Y. enterocolitica* of following adherence and invasion assay with ellipses drawn on average mean of each group.

### Arachidonic acid alters the peptide mass fingerprint of *Y. enterocolitica*

Following the cellular adhesion and invasion assay, Ye was plated on tryptic soy agar (TSA) to enumerate colonies. These same bacteria were also analyzed for changes in peptide mass fingerprints using matrix-assisted laser desorption/ionization – time of flight (MALDI-TOF) mass spectrometry (MS). MALDI-TOF MS has been useful for bacterial identification and has recently been validated in identifying strain-level and functional differences.^44^ Briefly, the MALDI-TOF MS creates a peptide mass fingerprint (PMF) through the ionization and excitation of the bacteria in a given matrix. The identification of bacteria to the strain level is achieved by comparing the PMFs to a curated database of known bacteria.^45^ We have previously tested the capacity of the MALDI-TOF MS to use PMFs to detect differences in the expression of protein caused by environmental factors, such as temperature, oxygen levels, and media.^46^ The MALDI-TOF MS was able to reliably detect differences in the expression of virulence factors between Ye grown at room temperature and 37°C.^46^ Using the MALDI-TOF MS, we compared the PMFs of Ye without exposure to epithelial cells, following exposure to epithelial cells, and intracellular Ye recovered after invasion. Principal component analysis of the peptide signatures from vehicle-treated Ye shows three different clusters, each corresponding to a different condition (without epithelial cells, with epithelial cells, and intracellular) (**Figure 3C**). These results demonstrate that Ye contains different protein signatures dependent upon the specific interaction with human epithelial cells. In contrast, this clustering is absent when Ye is treated with AA (Figure 3C). All three AA-conditioned Ye groups clustered together and appear to overlap with the intracellular Ye treated with vehicle control. Regardless of the presence of mammalian cells, Ye treated with AA expressed a protein signature similar to that of Ye found intracellularly. Taken together, these data suggest that treatment with AA induces adherence, invasion, and virulence products in Ye.

### *Y. enterocolitica* becomes more virulent *in vivo* after exposure to arachidonic acid

To determine if the AA-induced virulent state in Ye found *in vitro* could be replicated in the host, we tested AA-conditioned Ye in a mouse model of infection. Ye was pretreated with AA or vehicle control for 2 hours at 26°C and then washed prior to oral gavage in order to ensure that any residual AA was not administered to the mice. Inoculation of Ye was normalized for all mice to receive the same number of Ye (1×10^7^ CFU) (**Figure 4A**). Mice that received Ye conditioned with the vehicle did not exhibit weight loss in the first 3 days following infection (**Figure 4B**). In contrast, mice infected with AA-conditioned Ye rapidly lost a significant amount of weight beginning within the first two days after infection. Due to the visible discomfort and the rapid decline in health in the AA-conditioned group, the experiment was ended and all mice were euthanized at day 3 post-infection. Ye colonization of the ileum, Peyer’s patch, and spleen was assessed. Exposure to AA prior to infection did not increase the numbers of Ye in the ileum or Peyer’s patches compared to vehicle-treated Ye, though 100% of mice infected with the AA-conditioned Ye had colonization in the ileum compared to 50% of the mice receiving the vehicle-conditioned Ye (**Figure 4C**). Similar amounts of Ye was recovered in the Peyer’s patches regardless of Ye pre-treatment. Notably, none of the mice infected with vehicle-treated Ye had detectable Ye in the spleen while 66% of mice infected with the AA-conditioned Ye had colonization in the spleen. Ye typically causes weight loss beginning at 4 days post-infection with bacterial colonization of the spleen observed at day 6 (unpublished observations). These data indicate that exposure of Ye to AA prior to infection causes a more severe disease with rapid weight loss occurring just 48 hours after infection and dissemination of bacteria to the spleen.

**Figure 4.**
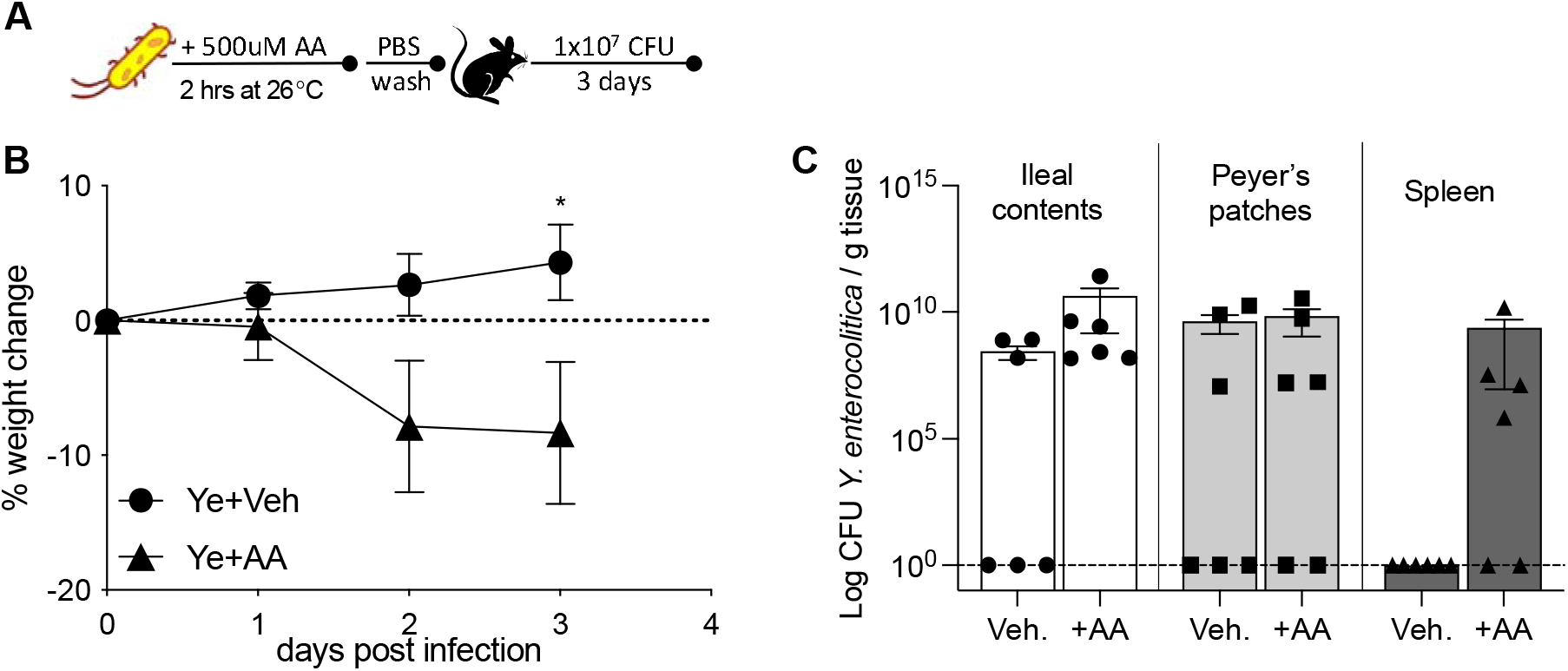
Exposure to arachidonic acid increases virulence Y. enterocolitica *in vivo*. **(A)** Diagram of mouse model with *Y. enterocolitica* pre-treated with 500 μM AA. **(B)** Percent weight change post infection. **(C)** Bacterial burden post infection at day 3 in the ileum, Peyer’s patches, and spleen. (B) Two-Way ANOVA with Bonferroni’s multiple comparisons test. Data is mean ± SEM from two independent experiments, n=6 per group, *P<0.05.

## DISCUSSION

Polyunsaturated fatty acids (PUFAs) are a vital component of our diet. As dietary recommendations are constantly changing, it is important to analyze the effects of dietary fats on potential bacterial food pathogens. In this study, we analyzed the effect of omega-6 PUFA arachidonic acid on *Y. enterocolitica*. We found that arachidonic acid significantly increases the growth of *Y. enterocolitica* and co-supplementation with eicosapentaenoic acid can suppress this growth in a dose-dependent manner. Supplementation with arachidonic acid also induces protein production much like the *Y. enterocolitica* grown at virulence-inducing temperatures. Indeed, our invasion assays demonstrated that *Y. enterocolitica* treated with arachidonic acid have more intracellular invasion and the MALDI-TOF MS analysis reveal peptide signatures similar to intracellular bacteria. Furthermore, we discovered that arachidonic acid-conditioned *Y. enterocolitica* has a more severe pathogenesis in the murine host. To our knowledge, this is the first study to show that a pathogen can have increased virulence in a host with altered pathogenesis after exposure to arachidonic acid.

There is a pronounced disconnect between dietary recommendations and the literature regarding the consumption of omega-6 (n-6) PUFAs. According to the 2015-2020 Dietary Guidelines for Americans provided by the U.S. Department of Health and Human Services, the recommended consumption of dietary oils should be 27 g per day for a 2,000 calorie diet.^47^ These guidelines do not distinguish between n-6 and n-3 PUFAs in the daily recommendations despite several studies finding that the excessive consumption of n-6 PUFAs in the American diet can be associated with inflammatory diseases. While n-6 PUFAs are necessary for proper development and growth, arachidonic acid has been positively correlated with heart disease,^48^ obesity, and diabetes.^49^ In rodent studies, diets high in n-6 fatty acids cause systemic low-grade inflammation and metabolic endotoxemia^50^ and increases the abundance of gut microbes associated with inflammation.^51^

More recently, diets high in n-6 PUFAs has also been associated with infections. Studies using *in vitro* cellular assays and mouse models have found that n-6 PUFAs may worsen inflammation when challenged with viral and bacterial pathogens.^52–54^ Whereas these studies use dietary models, our study introduces the bacterial pathogen to physiologically relevant levels of individual PUFAs prior to infection. Not only do we find the bacterial pathogen altered by arachidonic acid, the disease caused by the AA-treated *Y. enterocolitica* is worse in a murine host. Collectively, our study and others^37,55–57^ highlight the need to further investigate how pathogens interact with dietary components and how the consumption of diets excessive in fat may increase susceptibility to infections.

The literature reports that excessive dietary n-6 PUFAs is detrimental due to the increasing ratio of n-6 to n-3 PUFAs. As both n-6 and n-3 PUFAs rely on the same enzymes to breakdown into their respective metabolites, a healthy n-6:3 ratio is between 1:1 and 3:1 based on ancestral diets.^52–54^ Following the advent of the agricultural revolution, there has been an incline of n-6 PUFA consumption without a paralleling incline of n-3 consumption, leading to the Western diet having a PUFA ratio ranging from 10:1 to 20:1.^43,58^ Indeed, several human studies have shown that decreasing n-6:3 ratios can lower levels of inflammation associated with metabolic syndrome,^59^ obesity,^60^ non-alcoholic fatty liver disease,^61^ rheumatoid arthritis,^62^ and inflammatory bowel disease.^63^ With regards to infections, however, few studies examine how the n-6:3 ratio can affect bacterial pathogen susceptibility and none of those examine how the n-6:3 ratio can impact the pathogen. In our study, we find that co-exposure of arachidonic acid and eicosapentaenoic acid returns Ye proliferation to normal levels and prevents the increased invasion seen with arachidonic acid treatment alone. These studies identify a potential new role for n-3 PUFAs in the antagonism of n-6 induced bacterial virulence.

Studies have shown that arachidonic acid may also act as an antimicrobial agent. One study found that n-6, n-7, and n-9 fatty acids including arachidonic acid can inhibit the growth of oral bacteria.^37^ Another study found that arachidonic acid could inhibit the production of cholera toxin in *Vibrio cholerae*.^64^ Studies find long-chain fatty acids capable of inducing virulence as a potential signaling molecules. Golubeva and colleagues found host intestinal free long-chain fatty acids may also serve as a metabolic cue for *S. enterica* to start intestinal colonization.^56^ A recent study found that exposure to unsaturated fatty acids – palmitic and oleic acid – increased cytotoxin and flagella virulence genes in *Helicobacter pylori*.^65^ Further studies are needed to determine how other dietary factors can impact the virulence of *Y. enterocolitica*.

We found that arachidonic acid induces *Y. enterocolitica* induces the expression of virulence proteins in the absence of a temperature shift to 37°C and increases its invasiveness. Typically, *Y. enterocolitica* is considered an extracellular pathogen that interacts with the host cell surface, but recent studies have found and analyzed intracellular *Y. enterocolitica* populations.^66,67^. Arachidonic acid may act as an indicator to invade as we see increased intracellular invasion *in vitro* and which may also cause the quickening of disease *in vivo*.

To conclude, the present work demonstrates that PUFAs can differentially modulate *Y. enterocolitica* and affect its virulence. These types of studies provide valuable insight not only on the pathogen itself, but also how dietary factors may compromise the host.

## METHODS

### Bacterial strains and growth conditions

The strain used for this study is *Yersinia enterocolitica* 8081, biotype 1B serotype O:8. *Y. enterocolitica* was grown on tryptic soy agar (TSA) plates and in tryptic soy broth (TSB) for liquid cultures. For regular culturing, frozen stocks were plated on TSA and incubated at room temperature for 48 hours. After 48 hours, one colony forming unit was cultured in TSB and incubated overnight with agitation at room temperature. Subcultures were created by diluting overnight cultures 1:10 in TSB.

### Polyunsaturated fatty acid treatments and growth curves

Subcultures of *Y. enterocolitica* were treated with sterile phosphate-buffered saline (PBS), vehicle control, or 500 μM of the following polyunsaturated fatty acids unless otherwise stated: linoleic acid (LA), arachidonic acid (AA), alpha-linolenic acid (ALA) and eicosapentaenoic acid (EPA) (Cayman Chemicals, Ann Arbor, MI). PUFAs were diluted in 200 proof ethanol (VWR, Randor, PA) and were not used more than 3 openings to limit the amount of oxidation. *Y. enterocolitica* cultures were incubated at room temperature or 37°C with vigorous shaking and optical density (OD) at 600 nm was measured on a Genesys 30 Visible Spectrophotometer (ThermoFisher, Waltham, MA).

### SDS-PAGE and Coomassie Blue stain

*Y. enterocolitica* subcultures were supplemented with vehicle controls or PUFAs and incubated at room temperature or 37°C with agitation. After incubation for 2 hours, bacterial cells were separated by centrifugation at 4,000 x g for 10 min. Supernatant was collected and bacterial pellets were washed in PBS and resuspended in 100 μl of 10% sodium dodecyl sulfate (SDS) in PBS. Proteins from the supernatant and bacterial lysates were extracted by trichloroacetic acid (TCA) precipitation. Briefly, TCA was added to samples at a final concentration of 5% and incubated on ice for 2 hours. Protein was recovered by centrifuging at 13,000 x g for 10 minutes at 4°C and washed twice in ice-cold acetone. Samples were then resuspended in sample buffer and run on a 10% polyacrylamide pre-cast gel (Bio-Rad, Hercules, CA) for SDS-polyacrylamide gel electrophoresis (SDS-PAGE). The gel was stained with Coomassie blue stain containing 40% (v/v) methanol, 10% (v/v) glacial acetic acid, and 0.1% (w/v) Coomassie Brilliant Blue R-250 (ThermoFisher, Waltham MA) and de-stained until desired background was reached and imaged.

### Cellular adhesion and invasion assays

SW620s (ATCC; CCL-227) human intestinal epithelial adenoma cells were maintained in Dulbecco’s Modified Eagle Medium (DMEM) supplemented with 10% fetal bovine serum (FBS) and 200 μM L-glutamine. Prior to cellular assays, SW620s were seeded into 24-well tissue-culture treated plates and grown to 80-90% confluent monolayer over 3-4 days. Cells were rinsed with PBS and infected with *Y. enterocolitica* at a moiety of infection (MOI) of 10:1 with each starting dose plated on TSA to enumerate CFUs. Briefly, overnight *Y. enterocolitica* cultures were diluted 1:10 in TSB and supplemented with either 500 μM AA, equal volume ethanol as vehicle control, or PBS. Subcultures were then incubated for 2 hours at 26°C while shaking at 150 rpm. Cultures were washed in PBS, diluted to the appropriate infection dose in cell media (DMEM+10% FBS) and applied to SW620 cells that were previously rinsed 3 times with warm PBS. Infection was initiated by centrifuging the plates for 5 minutes at 500 x g and incubated at 37°C in a 5% CO_2_ incubator. For adhesion, *Y. enterocolitica* was co-cultured with SW620s for 1 hour. For intracellular invasion, *Y. enterocolitica* was co-cultured with SW620s for 1 hour, washed with PBS and then treated with 100 μg/ml gentamicin (Corning, Corning, NY) for an additional hour. *Y. enterocolitica* samples were eluted by washing the wells 3 times with warm PBS followed by treatment with 500 μl PBS with 1% Triton X-100 (AMRESCO, VWR, Randor, PA) for 5 minutes. Samples were then plated on TSA in serial dilutions to enumerate *Y. enterocolitica* colonization. Percentage of colonization was calculated by dividing the number of *Y. enterocolitica* CFUs recovered from co-cultures by the number of *Y. enterocolitica* CFUs initially applied to the wells.

### MALDI-TOF peptide-mass fingerprint analysis

*Y. enterocolitica* plated on TSA following the cellular and invasion assays were analyzed using matrix-assisted laser desorption/ionization – time of flight (MALDI-TOF) on Bruker’s Microflex LT/SH (Bruker, Billerica, MA). Protein was extracted per manufacturer’s protein extraction method.^68^ Briefly, CFUs were picked into sterile water, homogenized, and treated with 100% ethanol (Fisher Chemical). After centrifuging and air-drying the protein pellets, the protein pellets were resuspended in equal volumes 70% formic acid (Fisher Chemical, Hampton, NH) and acetonitrile (Fisher Chemical). The protein extracts were centrifuged and 1 μl of supernatant was plated on stainless-steel target in triplicates. Samples were then overlaid with 1 μl of alpha-Cyano-r-hydroxycinnamic acid (HCCA) Matrix (Bruker, Billerica, MA) and analyzed using the Microflex (Bruker). For MALDI-TOF standard, Bruker’s bacterial test standard (BTS) (Bruker) was used.

Raw spectra text files were analyzed using the R package MALDIquant [https://www.ncbi.nlm.nih.gov/pubmed/22796955]. Raw spectra data were trimmed to a spectrum range of 3,000 to 15,000 m/z. The spectral intensities were then square-root transformed and smoothed using the Savitzky-Golay algorithm. Baseline noise was removed using the statistics-sensitive non-linear iterative peak clipping (or SNIP) algorithm with 100 iterations. The data were then normalized using total ion current (or TIC) calibration, which sets the total intensity to 1. Multiple spectra within the same analysis were aligned to the same x-axis using the Lowess warping method, a signal-to-noise ratio of 3, and a tolerance of 0.001. Peaks were detected from the average of 4 technical replicates using median absolute deviation. Principal components analyses and hierarchical clustering were also performed in R using the base statistics package. Hierarchical clustering was performed on a calculated Euclidean distance matrix using Ward’s method.

### Oral mouse infections

C57Bl/6 male mice 4-5 weeks of age were used for *in vivo* infections. *Y. enterocolitica* treated with either vehicle control or 500 μM AA was washed and resuspended in PBS. Mice were given oral infection of 100 μl of 1×10^8^ CFU/ml using a sterile blunt-ended needle and infection was carried out for 3 days. Mice weight and fecal consistency was monitored daily. After 3 days, mice were euthanized and tissue was collected. Ileal contents, Peyer’s patches, and spleens were aseptically extracted and plated in dilutes on *Y. enterocolitica* selective plates (Fisher, Waltham MA) or TSA. *Y. enterocolitica* CFUs were enumerated after 48 hours. All animal experiments were performed at the University of Washington, Seattle following experimental review and approval by the Institutional Biosafety Committee and the Institutional Animal Care and Use Committee.

### Statistical analysis

All statistics were performed on GraphPad Prism v7 (GraphPad Software, San Diego, CA, www.graphpad.com) unless stated otherwise. One-way or two-way ANOVA tests were used to determine the significance of the experiments as indicated in each figure legend. Data is shown as mean with bars showing standard error of the mean.

## ACKNOWLEDGEMENTS

The authors thank Leandra Brettner for her help in analyzing the MALDI-TOF MS data.

## SUPPLEMENTAL FIGURE

**Supplemental Figure 1.**
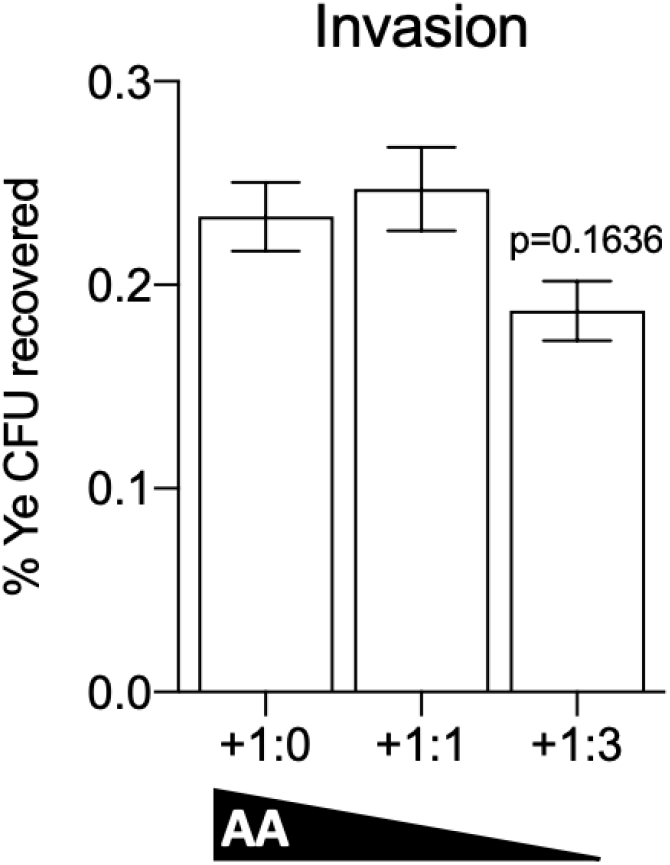
Percentage *of Y. enterocolitica* CFUs recovered after 60 minutes of incubation then treatment of 100 μg/ml gentamicin for 60 minutes. One-way ANOVA with Tukey’s multiple comparisons test. Data is mean ± SEM from a representative from 2 independent studies with duplicates.

